# Unstructured regions differentially modulate the activation of RBOHD and RBOHH

**DOI:** 10.64898/2026.03.31.715192

**Authors:** Matteo Citterico, Michaela Neubergerová, Alexis Porcher, Adam Zeiner, Přemysl Pejchar, Tereza Korec Podmanicka, Martin Potocký, Saijaliisa Kangasjärvi, Roman Pleskot, Michael Wrzaczek

## Abstract

Reactive oxygen species (ROS) produced by plant NADPH oxidases (RBOHs) must be precisely controlled to generate appropriate signaling outputs in profoundly diverse physiological contexts. RBOHs are activated by Ca²⁺ and phosphorylation, yet how structural variation in their regulatory regions shapes these responses and contributes to functional specialization remains unclear. To identify regions underlying functional diversification among RBOHs, we performed phylogenetic and sequence-comparison analyses across land plants. The most variable regions are two N-terminal unstructured regions (URs), UR1 and UR2, which flank the EF-hand Ca²⁺-binding domain (EFD). We investigated whether these unstructured regions contribute differently to the regulation of Arabidopsis RBOHD and RBOHH, which function predominantly in immune signaling and pollen tube growth, respectively. Our functional assays in HEK293T cells suggest that UR1 plays opposing roles in RBOHD and RBOHH, restraining Ca²⁺-dependent activation in RBOHD while facilitating it in RBOHH. Our analyses indicate differences in the regulatory contribution of UR2 between RBOHD and RBOHH. In RBOHD, our results suggest that UR2 phosphorylation promotes formation of an α-helical conformation and its interaction with the catalytic domain, contributing to enzymatic activation, whereas our analyses do not support a comparable regulatory mechanism in RBOHH. Domain-swapping experiments between RBOHD and RBOHH suggest that the EFD coevolved with UR2 in RBOHD to enhance phosphorylation-dependent activation, whereas in RBOHH it coevolved with UR1 to favor Ca²⁺-dependent activation. Together, our findings support a model in which diversification of unstructured regulatory regions adapts the conserved RBOH enzymatic core to distinct physiological contexts, including immune signaling and polarized growth.

## Introduction

Reactive oxygen species (ROS) play dual roles in plants, acting both as damaging oxidants and essential signaling molecules that orchestrate plant growth, development, and stress responses (Castro *et al*, 2021; Zhang *et al*, 2025; Waszczak *et al*, 2018). Precisely regulated ROS production is fundamental to maintaining cellular homeostasis and enabling rapid responses to environmental challenges. In land plants, stimulus-triggered extracellular ROS bursts are predominantly generated by plasma membrane-localized NADPH oxidases known as Respiratory Burst Oxidase Homologues (RBOHs) (Zhang *et al*, 2023; Cheng *et al*, 2020; Liu *et al*, 2020). These enzymes integrate pathogen-derived signals, abiotic stresses, developmental inputs, and ion fluxes to generate a controlled ROS production in response to specific physiological demands. Accordingly, RBOH-driven ROS production is crucial for pathogen defense, abiotic stress responses, and developmental processes such as stomatal movement, root growth, and pollen tube elongation (Torres *et al*, 2002; Lee *et al*, 2020; Ngou *et al*, 2021; Arnaud *et al*, 2023; Orman-Ligeza *et al*, 2016; Kaya *et al*, 2014). RBOH activation depends on two major regulatory inputs: Ca²⁺ binding to an N-terminal EF-hand domain (EFD) and phosphorylation by protein kinases (Hashimoto *et al*, 2023; Ogasawara *et al*, 2008). These inputs integrate environmental and developmental cues to tune ROS production with high spatio-temporal precision.

To ensure appropriate control of ROS production across diverse physiological contexts, *Arabidopsis thaliana* encodes ten RBOH proteins (RBOHA–RBOHJ). Among them, RBOHD is broadly expressed with central roles in both stress-related and developmental processes (Chen & Yang, 2020; Kadota *et al*, 2015), whereas RBOHH is specifically expressed at the pollen tube tip, where its activity drives tube elongation and prevents bursting (Kaya *et al*, 2014). Comparative studies in human embryonic kidney (HEK) 293T cells have shown that RBOHH produces roughly tenfold higher ROS levels than RBOHD in response to elevated cytosolic Ca²⁺, consistent with the strong Ca²⁺ oscillations that underlie pollen tube growth (Kaya *et al*, 2019). By contrast, RBOHD is a well-established phosphorylation target for several protein kinases, including BOTRYTIS-INDUCED KINASE 1 (BIK1) (Li *et al*, 2014; Kadota *et al*, 2014), calcium-dependent protein kinases (CPKs) (Köster *et al*, 2025; Dubiella *et al*, 2013; Yu *et al*, 2024), and CYSTEINE-RICH RECEPTOR-LIKE KINASE 2 (CRK2) (Kimura *et al*, 2020). Phosphorylation-dependent activation is essential for RBOHD function in immunity: mutation of three key serine residues (S339, S343, S347) to non-phosphorylatable alanine almost completely abolishes flg22-triggered ROS production *in planta* (Li *et al*, 2021), even though flg22 perception triggers robust Ca²⁺ influx and broad phosphorylation cascades that converge on multiple sites within RBOHD besides S339, S343 and S347 (Li *et al*, 2014; Kadota *et al*, 2014, 2019).

Yet, despite the clear evidence that RBOHH has a higher capacity for Ca²⁺-induced activation compared to RBOHD, the molecular basis for this difference remains unclear. Likewise, although phosphorylation of S339, S343, and S347 is essential for full RBOHD activation, it is not understood how these phosphorylation events are translated into catalytic activation.

Here, we investigated these two modes of RBOH activation separately. First, we examined the molecular basis for the stronger Ca²⁺-induced activity of RBOHH compared with RBOHD. Through phylogenetic and AlphaFold-based structural analyses, we identified UR1, the most N-terminal unstructured region of RBOHs, as the most variable region across the family. Using HEK293T cells, which lack plant-specific signaling components such as CPKs, CIPKs, and HPCA1-mediated signaling, we examined how RBOHD and RBOHH respond directly to elevated cytosolic Ca²⁺. This system allowed us to assess the contribution of UR1 to Ca²⁺-dependent RBOH activation independently of these plant-specific regulatory pathways. Our results suggest that divergence of UR1 is associated with opposite regulatory roles in RBOHD and RBOHH. Using chimeric proteins, we further found that UR1 and the EFD in RBOHH appear to have co-evolved to enhance Ca²⁺-induced activation. Second, we investigated the molecular basis of phosphorylation-mediated activation of RBOHD. By combining molecular dynamics simulations, structural modeling, mutagenesis, and HEK293T-based activity assays, we propose a mechanism by which phosphorylation of S339, S343, and S347 activates RBOHD. Our data suggest that phosphorylation of these residues stabilizes an α-helical conformation within UR2, the second highly variable unstructured region of RBOHs. This phosphorylated UR2 helix promotes electrostatic interactions with positively charged residues in the catalytic core, thereby strengthening the interaction between the N-terminal regulatory region and the C-terminal catalytic domain. In contrast, the corresponding UR2 region of RBOHH does not appear to undergo the same phosphorylation-dependent stabilization, suggesting that this activation mechanism is specific to RBOHD.

Thus, our findings indicate that RBOHH and RBOHD rely on distinct regulatory strategies. RBOHH is oriented toward strong Ca²⁺-induced activation through functional coupling between UR1 and the EFD, whereas RBOHD is oriented toward phosphorylation-induced activation through coupling between UR2 and the catalytic core. Together, these results show how divergent unstructured regions tune a conserved RBOH catalytic core toward different activation modes and physiological functions.

## Materials and Methods

### Plasmid Construction and Generation of Transgenic Lines

Deletion constructs of RBOHD and RBOHH and point mutants in the α-2 helix were generated using a modified QuickChange protocol optimized for Phusion DNA polymerase using partially overlapping primers (Xia *et al*., 2015). Templates were pcDNA3.1 vectors encoding 3×FLAG– RBOHD and 3×FLAG–RBOHH (Kimura *et al*., 2020). RBOHD–RBOHH chimeras in pcDNA3.1 were generated using PIPE cloning (Klock & Lesley, 2009).

The following Arabidopsis thaliana lines, all in the Columbia-0 (Col-0) background, were used in this study: *35S–RBOHD/rbohD* (Kimura *et al*., 2020), the *rbohd* knockout mutant (Torres *et al*., 2002), and *35S–*χ*3/rbohD*. The χ3 construct was generated by replacing the RBOHD coding sequence in pBm43GW 35S–RBOHD with the χ3 sequence using PIPE cloning. The construct was introduced into the *rbohd* background by *Agrobacterium tumefaciens* GV3101 (pSoup)-mediated floral dip. Plants were grown in a 1:1 mixture of peat and vermiculite under a 12 h light/12 h dark photoperiod, at 22°C during the day and 18°C at night, with a light intensity of 200 µmol m⁻² s⁻¹ and relative humidity of 60% during the day and 70% at night.

### Cell Culture and Transfection

HEK293T cells (ATCC, CRL-3216) were maintained at 37 °C in 5% (v/v) CO₂ in Dulbecco’s Modified Eagle’s Medium nutrient mixture Ham’s F-12 (Sigma D8062), supplemented with 10% (v/v) fetal bovine serum (Gibco, 26140-079). For ROS assays, cells were diluted to 1.5 × 10^5^ cells mL^−1^, and 130 µL of the suspension was seeded into each well of white poly-D-lysine-coated 96-well plates (Corning, 356620). Cells were incubated for 24 h before transfection. For transfections, each well received 120 ng plasmid DNA mixed with 6.3 µL Opti-MEM (Life Technologies, 31985-062) and 0.36 µL GeneJuice transfection reagent (Merck Millipore, 70967-3). For co-transfections, a total of 120 ng DNA was used; for example, RBOHD and BIK1 were transfected at 100 ng and 20 ng per well, respectively. Transfected cells were incubated for 48 h at 37°C and 5% CO_2_ before ROS measurements.

### Protein extraction and Immunoblotting

Plant tissue was frozen in liquid nitrogen and ground to a fine powder before protein extraction using 50 mM Tris-HCl (pH 7.5) containing 2% SDS. HEK293T cells were lysed directly in the wells using 75 µL of 1x Laemmli buffer supplemented with 300 mM DTT. Lysates were incubated at room temperature for 10 min without heating to avoid RBOH aggregation. Protein expression in plant tissue and HEK293T cells was analyzed by immunoblotting using an anti-FLAG antibody (Sigma, F1804), followed by IRDye800CW anti-mouse IgG (LI-COR, 926-32210). Equal loading was verified using anti-β-actin (Sigma, A5316) for HEK293T samples or by Coomassie staining for plant samples.

### ROS Measurements and Statistical Analysis

Two days after transfection, HEK293T cells were washed with HBSS (Gibco, 14025-092). Measurements were initiated 1 min after addition of HBSS containing 250 µM luminol and 20 mg/L horseradish peroxidase. Ionomycin (1 µM; Calbiochem, 407952) was added to induce Ca²⁺ influx. Luminescence was recorded at 37 °C using a Tecan Spark luminometer, and values are presented as mean ± SE. ROS signals were normalized to the maximum mean response of the reference construct within each experiment. ROS assays were performed in three independent experiments, each with five technical replicates per condition. Total ROS production was quantified as the area under the curve (AUC) using trapezoidal integration. Statistical comparisons were performed on AUC values using one-way ANOVA followed by Tukey’s HSD test (different letters indicate statistical difference, *P* < 0.05).

In Arabidopsis, 4mm leaf discs from 4-week-old plants were floated overnight in distilled water under continuous light. Measurements were initiated 1 min after addition of assay buffer containing 250 µM luminol, 20 mg/L horseradish peroxidase, and 200 nM flg22 (GenScript). ROS signals were analyzed as described for the HEK293T cell assays.

### Phylogenetic and Sequence Analyses

RBOHD, RBOHH, and FRO1 protein sequences were used to identify homologs using BLASTP and TBLASTN searches in NCBI, Phytozome, and Phycocosm databases. Additional queries included human NOX2, NOX5, DUOX1, and yeast FRE1. Sequences were aligned using MAFFT E-INS-I (Katoh & Standley, 2013) implemented in Jalview (Waterhouse *et al*., 2009), and gap-rich and poorly conserved regions were removed. Phylogenetic inference was performed using IQ-TREE with ultrafast bootstrap support (Nguyen *et al*., 2015), and Bayesian phylogenetic inference of the embryophyte RBOH dataset was performed using MrBayes (Ronquist *et al*., 2012). Detailed phylogenetic parameters are provided in the Supplementary Methods. Domain boundaries (UR1, UR1cts, EFD, UR2, catalytic core, helix α-2) were defined on the AlphaFold3 model of Arabidopsis RBOHD and mapped onto the trimmed embryophyte RBOH alignment as fixed column intervals (e.g. UR1, alignment columns 1–549, corresponding to residues 1–170 of RBOHD). Analyses were restricted to the eudicot RBOH subset (232 sequences, Supplementary Dataset S5). Within each subfamily, every sequence was compared with the Arabidopsis reference protein of that subfamily (RBOHA, RBOHB, RBOHD, RBOHE, RBOHF and RBOHH for RbohA/C/G, B, D, E, F/I and H/J, respectively) using the existing multiple-sequence alignment. Domain length was taken as the number of non-gap residues of the query within the domain interval. Sequence identity was calculated as the number of alignment columns at which query and reference carry the same residue, divided by the number of columns at which at least one of the two carries a residue. Distributions were compared across subfamilies by Kruskal–Wallis test followed by pairwise two-sided Mann–Whitney U tests with Benjamini–Hochberg correction. Sequence datasets and alignments are provided as Supplementary Datasets S1–S5.

### Structural Modelling and Molecular Dynamics Simulations

Structures of phosphorylated and non-phosphorylated *Arabidopsis thaliana* RBOHD and RBOHH were predicted using the AlphaFold3 web server (Abramson *et al*., 2024). Molecular dynamics simulations were performed using GROMACS version 2020.3 (Abraham *et al*., 2015). Systems were prepared using CHARMM-GUI v3.8 (Jo *et al*., 2008; Wu *et al*., 2014) with the CHARMM36m all-atom force field (Huang *et al*., 2017). Simulations were performed for phosphorylated and non-phosphorylated RBOHD and RBOHH UR2 regions in water and for phosphorylated and non-phosphorylated RBOHD171–921 embedded in a model plant plasma membrane. Detailed simulation setup, equilibration, production parameters, and analyses are provided in the Supplementary Methods.

## Results

### Structural and Evolutionary Analysis of calcium-binding domain and unstructured regions

To examine how functional domains have diversified within the RBOH family, we first analyzed structural models of the ten Arabidopsis thaliana RBOH proteins predicted by AlphaFold3, together with sequence alignments. Generally, RBOHs share a highly conserved catalytic core module composed of a transmembrane region formed by six helices that coordinate two heme groups, followed by a C-terminal cytosolic dehydrogenase domain containing the FAD- and NADPH-binding sites. This core module is predicted with high structural confidence (Fig. S1A,B) and constitutes the electron transport chain that transfers electrons from cytosolic NADPH via FAD to extracellular oxygen, producing superoxide (Fig. 1A).

**Figure 1.**
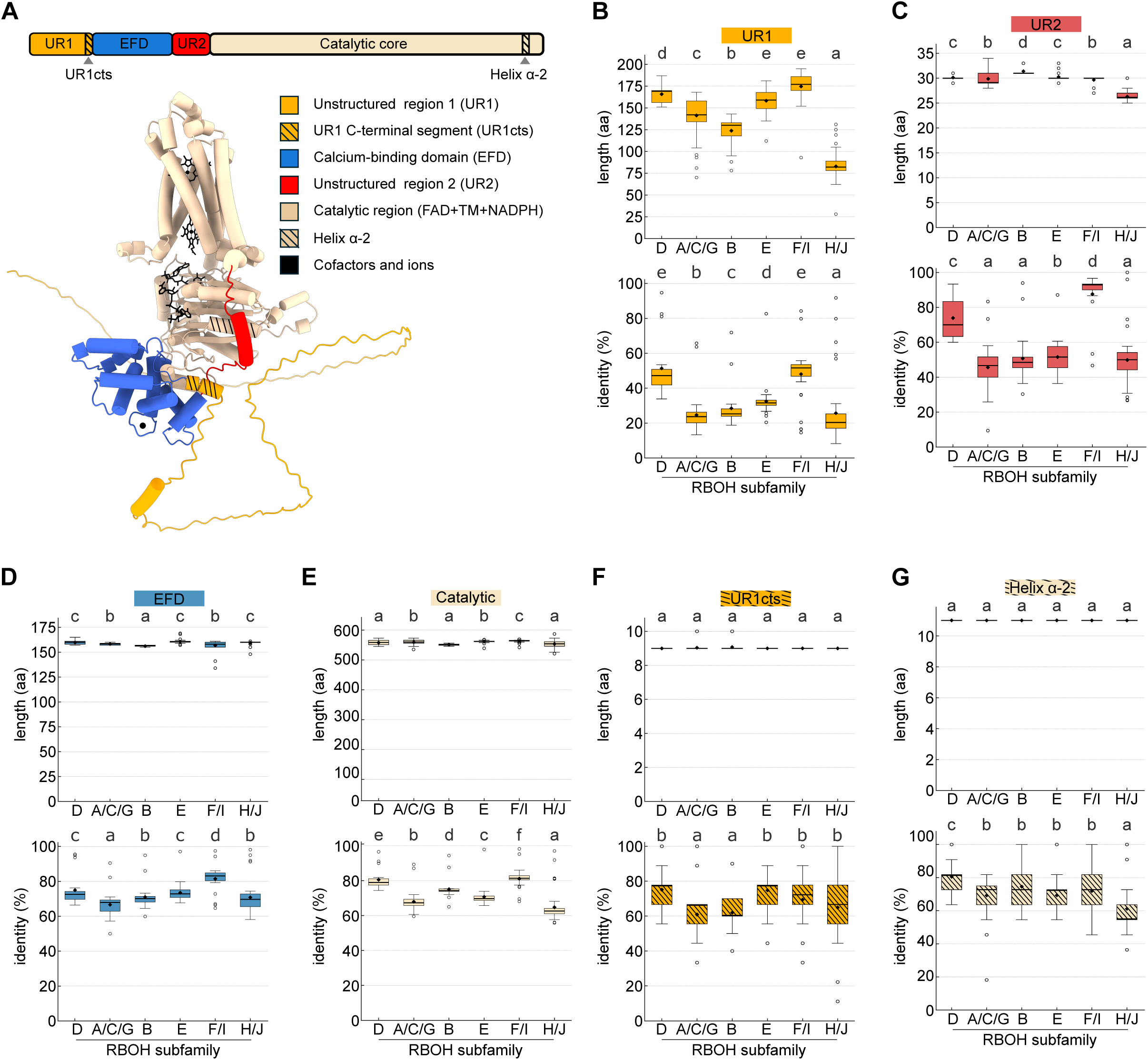
Structural organization and evolutionary diversification of RBOH protein subdomains. **(A)** Linear representation of RBOH structure together (upper part) with the AlphaFold-predicted structure of Arabidopsis RBOHD (lower part). Helices are shown as tubes; protein domains and cofactors are colored as indicated in the figure. **(B-G)** Length and sequence-conservation profiles of RBOH structural subdomains across the six RBOH subfamilies in eudicots. Different letters indicate statistically significant differences between subfamilies (Kruskal–Wallis test with Mann–Whitney post hoc comparison with Benjamini–Hochberg FDR correction, P < 0.05).

In addition to the catalytic module, RBOHs possess a cytosolic N-terminal regulatory domain composed of a structured EFD flanked by two intrinsically disordered regions: Unstructured region 1 (UR1) at the N-terminus and unstructured region 2 (UR2), which connects the EFD to the transmembrane region (Fig. 1A). The EFD consists of α-helical bundles connected by short loops, forming a compact but flexible domain that is predicted with high confidence by Alphafold3 (Fig. S1A,B). Two canonical EF-hand Ca²⁺-binding motifs are located toward its C-terminal side, and Ca²⁺ binding is thought to trigger a repositioning of the α-helices of the EFD in a way that promotes catalytic activation (Gifford *et al*, 2007). Compared to the EFD, the structures of UR1 and UR2 are mostly predicted with lower confidence (Fig. S1A,B) and show high sequence divergence among the ten Arabidopsis RBOH paralogs (Fig. S2A,B). In Arabidopsis RBOHs, UR1 varies markedly in length, ranging from 88 amino acids in RBOHB to 181 in RBOHF. In RBOHD, UR1 is 170 amino acids long, whereas in RBOHH it comprises 109 residues, and the two share only 35–40 % sequence similarity (Fig. S2A). The predicted structure and topological position of UR1 relative to the catalytic core vary substantially across different AlphaFold3 models of the same RBOH, resulting in a high predicted alignment error (PAE) (Fig. S1C,D). This lack of reliability suggests that UR1 is highly disordered and may not maintain a stable spatial relationship with the structured portion of the protein. UR2 shows a slightly higher degree of predicted structural confidence compared to UR1 (Fig. S1A,B), and its length is relatively consistent across the Arabidopsis RBOH family (27–33 amino acids) (Fig. S2C). Although sequence similarity between the UR2 regions of RBOHD and RBOHH is low (25–30%), AlphaFold models consistently predict UR2 in a defined topological position relative to the catalytic core, adjacent to the penultimate helix of RBOHs (α-2), which forms part of the NADPH-binding pocket (Fig. 1A).

To assess domain-specific diversification across RBOH subfamilies, we compared the length and sequence conservation of UR1, the EFD, UR2, and the catalytic core across major eudicot RBOH subfamilies (Fig. 1B–F). These subfamilies were defined through a focused phylogenetic analysis of 479 RBOHs from 131 embryophyte species (Fig. S3), which resolved seven well-supported eudicot RBOH subfamilies, six containing Arabidopsis RBOH isoforms: RbohA/C/G, RbohB, RbohD, RbohE, RbohF/I, and RbohH/J. Consistent with the patterns observed among Arabidopsis paralogs, the EFD and catalytic core showed relatively little variation and were strongly conserved, whereas the unstructured regions were considerably more variable (Fig. 1B– F). UR1 emerged as the most divergent region across subfamilies, both in length and sequence identity (Fig. 1B). RbohD and RbohF/I possess the longest and most conserved UR1 regions, with median lengths and sequence identities of 169 aa and 47% for RbohD and 177 aa and 52% for RbohF/I, respectively. In contrast, most RbohH/J orthologs contain short and highly variable UR1 regions (median length 82 aa; median sequence identity 20%). A similar, although less pronounced, pattern was observed for UR2 (Fig. 1C). By contrast, the EFD is highly consistent in length across subfamilies (median length ∼160 aa) and strongly conserved, with median sequence identities of 73% for RbohD and 70% for RbohH/J (Fig. 1D). Similarly, the catalytic core showed little variation in length and was highly conserved across RBOH subfamilies (Fig. 1E).

The RBOH phylogeny in embryophytes also revealed broader patterns in the evolutionary distribution of these RBOH subfamilies. Bryophyte orthologs remained positioned near the base of the RBOH clade, whereas lycophytes and ferns showed a clear separation of RBOH orthologs into distinct subfamilies (Fig. 2A; Fig. S3). The RbohH/J lineage was already distinct in lycophytes, whereas the RbohF/I and RbohE subfamilies were established in gymnosperms. The RbohA/C/G subfamily was reliably identified in eudicots, whereas the RbohD lineage was restricted to rosids (Fig. S3). Interestingly, our analysis also identified a previously unrecognized early-diverging RBOH subfamily present in lycophytes, ferns, gymnosperms, monocots, and most eudicots but absent from Arabidopsis (Fig. 2A).

**Figure 2.**
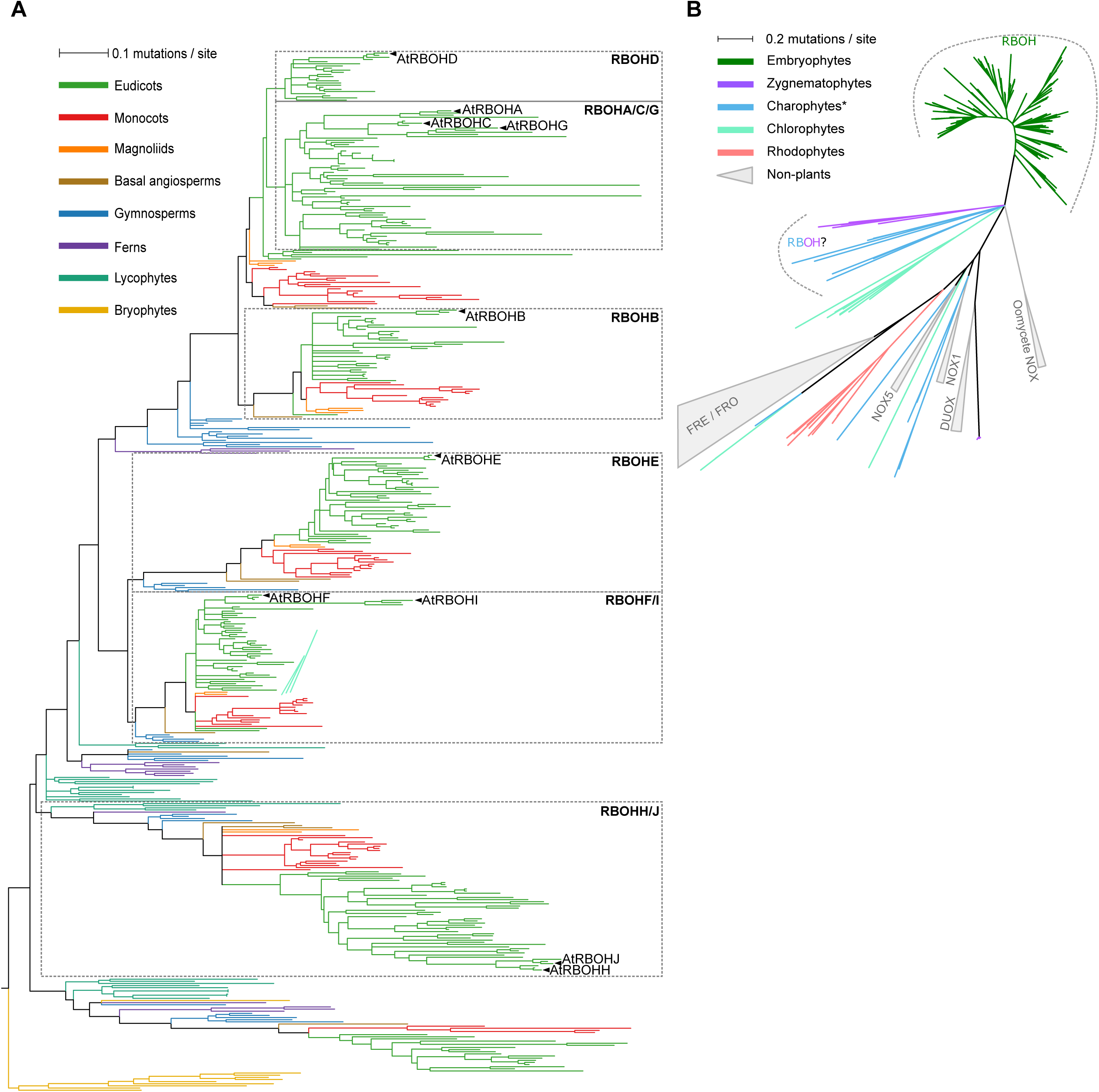
Evolution of the RBOH family. **(A)** High-resolution phylogeny of 479 embryophyte RBOHs from 131 species, showing the emergence and relationships among six major RBOH subfamilies (RbohA/C/G, B, D, E, F/I, and H/J). The color code denotes distinct plant groups, and Arabidopsis RBOH isoforms are highlighted. The full tree is shown in SI Appendix, Fig. S3. **(B)** Global schematic phylogeny of 304 bona fide NADPH oxidase– and ferric-reductase–related proteins across major eukaryotic lineages. The color code denotes major Archaeplastida lineages. DUOX, dual oxidase; FRE/FRO, ferric reductase/oxidase; NOX, NADPH oxidase; RBOH, respiratory burst oxidase homolog. See Supplementary Fig. S4 for the detailed phylogenetic tree.

We next asked when the characteristic RBOH architecture, including the paired EF-hand motifs, emerged during plant evolution. We therefore reconstructed a phylogeny of NADPH oxidase-related enzymes from representative algal and plant genomes and compared them with ferric reductase/oxidases (FRE/FRO), a related protein family (Fig. 2B; Fig. S4). Our analysis showed that canonical plant RBOHs with sequentially conserved EF-hand motifs and a conserved but structurally flexible N-terminal extension are already present in streptophyte algae such as Zygnematophyceae and Coleochaetales, suggesting that they evolved prior to the colonization of land. These *bona fide* RBOHs are clearly distinct from non-plant EF-hand-bearing NOX5, NOX1– 4 and DUOX subfamilies, as well as from ferric reductases (FRE/FRO). Our results suggest that the acquisition of paired EF-hand motifs, potentially independently of NOX5, represented a defining RBOH innovation during streptophyte evolution.

Taken together, these analyses suggest that the characteristic RBOH architecture emerged early in streptophyte evolution, whereas its regulatory regions subsequently diversified among land-plant RBOH lineages. In particular, UR1 and UR2 show extensive diversification across eudicot RBOH subfamilies, while the EFD and catalytic core remain strongly conserved. Among these subfamilies, RbohD shows relatively high conservation of its regulatory regions, whereas RbohH/J is considerably more variable.

### UR1 limits Ca²^⁺^-induced RBOHD activation in HEK293T cells

In RBOHD, UR1 (residues 1–170) has previously been reported to undergo phosphorylation at multiple residues, resulting in enzymatic activation and subsequent ROS production. However, whether UR1 also contributes to RBOHD regulation independently of these phosphorylation events remains unclear. To examine the contribution of UR1 to Ca²⁺-dependent RBOHD activation, we used HEK293T cells, which provide a heterologous system lacking plant-specific signaling pathways and therefore allow the response of RBOHD to elevated cytosolic Ca²⁺ to be assessed independently. To investigate the impact of UR1 on RBOHD activity in the non-phosphorylated state, we generated constructs encoding truncated variants of 3x-FLAG-RBOHD lacking portions of UR1 (DΔ60, DΔ69, DΔ83, DΔ100, DΔ121, DΔ134, DΔ155, DΔ170) and EFD (DΔ181, DΔ253) and transiently expressed them in HEK293T cells. Comparable expression of all constructs was confirmed by immunoblotting (Fig. S5A,B). ROS production was measured before and after treatment with ionomycin, which triggers Ca²⁺ influx in HEK cells and consequently activates RBOHD through Ca²⁺ binding to the EFD. Following ionomycin treatment, the DΔ60 and DΔ69 variants displayed ROS-producing activity comparable to that of full-length RBOHD (Fig. 3A). By contrast, DΔ83, DΔ100, and DΔ121 exhibited a progressive reduction in activity, with DΔ121 showing the strongest impairment (Fig. 3A). Unexpectedly, DΔ134 and DΔ155 restored activity to wild-type levels, while complete removal of UR1 (DΔ170) resulted in elevated ROS production under basal conditions and following ionomycin treatment (Fig. 3B). Additionally, partial loss of EFD in DΔ181 and DΔ253 completely abolished the ROS-producing activity (Fig. 3B).

**Figure 3.**
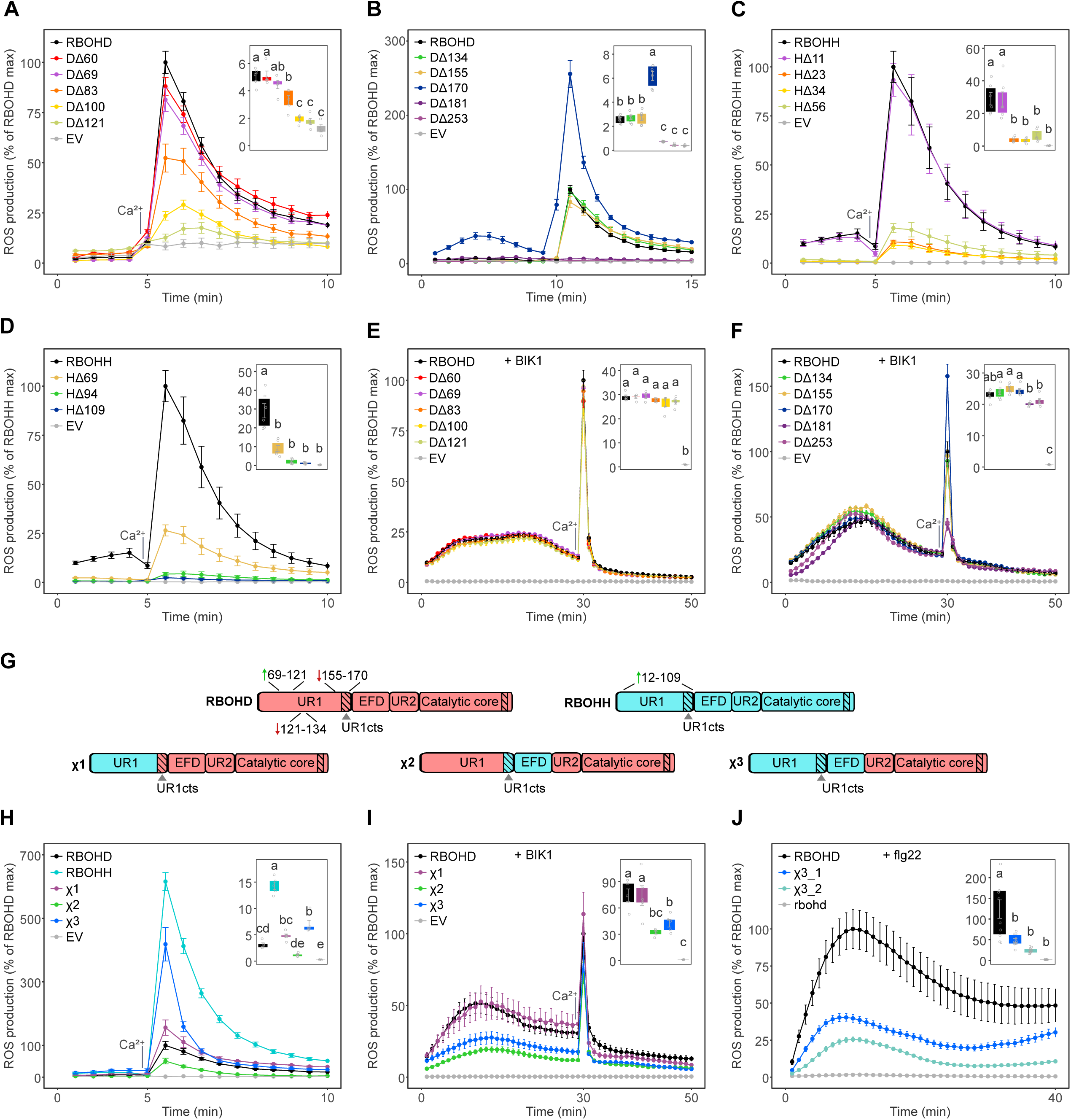
ROS production by truncated and chimeric RBOHD and RBOHH. (A,B) Ca²⁺-induced ROS production in HEK293T cells expressing 3×FLAG–RBOHD variants carrying progressive deletions in the UR1 region (DΔ60, DΔ69, DΔ83, DΔ100, DΔ121, DΔ134, DΔ155, DΔ170) or in the EF-hand domain (DΔ181, DΔ253). **(C,D)** Ca²⁺-induced ROS production in HEK293T cells expressing 3×FLAG–RBOHH variants with deletions in the UR1 region (HΔ11, HΔ23, HΔ34, HΔ56, HΔ69, HΔ94, HΔ109). **(E,F)** BIK1-induced and BIK1 + Ca²⁺-induced ROS production in HEK293T cells expressing RBOHD UR1 truncation variants. **(G)** Schematic representation of chimeric RBOHD–RBOHH variants (χ1, χ2, χ3). Domains derived from RBOHD are shown in pink and domains derived from RBOHH in cyan. **(H)** Ca²⁺-induced ROS production by chimeric RBOHD–RBOHH variants. **(I)** BIK1-induced and BIK1 + Ca²⁺-induced ROS production in HEK293T cells expressing chimeric RBOHD–RBOHH variants. **(J)** ROS production in leaf discs from plants overexpressing RBOHD or the χ3 chimera following flg22 treatment. ROS production is expressed as a percentage of the maximal response of the corresponding wild-type reference construct; in panel H, RBOHD was used as the reference. For Ca²⁺-induced conditions (A–D,H), luminescence was recorded every 1 min before ionomycin addition (indicated in the graphs) and every 30 s thereafter. For BIK1-dependent conditions (E,F,I) and flg22 treatment (J), luminescence was recorded every 1 min before and after stimulation. Data show one representative experiment of three independent experiments, with five technical replicates per condition (mean ± SEM). Inset shows the area under the ROS production curves (AUC), with y-axis values expressed in units of 10. AUC values were compared using one-way ANOVA followed by Tukey’s HSD test. Groups sharing no letters differ significantly (*P* < 0.05).

Taken together, these results suggest that UR1 (residues 1–170) functions as an autoinhibitory module in RBOHD, since its complete removal (DΔ170) enhanced Ca²⁺-induced ROS production beyond wild-type levels. The main inhibitory effect was exerted by the terminal segment (155– 170), directly adjacent to the EFD, as its deletion increased activity above that of the full-length protein. Our truncation analysis also revealed that UR1 might contain internal regulatory elements upstream of this primary inhibitory segment. Specifically, deletion of residues 69–121 reduced the Ca²⁺-induced activity of RBOHD. However, this loss was rescued when the downstream region spanning residues 121–134 was subsequently removed. This indicates that residues 121–134 could also function as an autoinhibitory segment, and that its inhibitory effect can only be effectively relieved in the presence of the upstream 69–121 region.

### UR1 is required for Ca²^⁺^-induced RBOHH activation in HEK293T cells

Because UR1 differs substantially between RBOHD and RBOHH in both length and amino acid composition, we investigated whether these differences could result in a distinct regulatory role of the given UR1 on enzymatic activity. We generated a series of truncated variants of RBOHH (HΔ11, HΔ23, HΔ34, HΔ56, HΔ69, HΔ94, and HΔ109) and transiently expressed them in HEK293T cells to isolate their Ca²⁺-induced activity. All constructs were expressed at comparable levels (Fig. S5C). ROS production was quantified before and after ionomycin treatment.

The HΔ11 variant of RBOHH exhibited ROS production comparable to the full-length protein (Fig. 3C). A noticeable reduction in activity was observed with HΔ23 (Fig. 3C). The HΔ34 and HΔ56 truncations of RBOHH displayed even more reduced ROS-producing activity (Fig. 3C). HΔ69 displayed a slight increase over HΔ56, while HΔ94 and HΔ109 nearly abolished the ROS-producing activity (Fig. 3D).

Taken together, these results suggest that, in contrast to RBOHD, UR1 positively contributes to Ca²⁺-induced activation of RBOHH in HEK293T cells, with progressive removal of the region strongly impairing ROS production. The opposite effects of UR1 truncation in RBOHD and RBOHH were particularly evident near the C-terminal end of UR1, immediately upstream of the EFD. In RBOHD, removing residues 155–170 increased Ca²⁺-induced activity, whereas deleting the corresponding region in RBOHH (residues 94–109) strongly impaired ROS production. Sequence comparison showed that the proximal part of this region is relatively conserved between RBOHD and RBOHH. The most C-terminal residues are also relatively conserved across the RbohD and RbohH/J clades, but show several non-conservative substitutions between Arabidopsis RBOHD and RBOHH, including an HA dipeptide in RBOHD that is replaced by RG in RBOHH/J (Fig. 1F; Fig. S2C). In the subsequent analyses, we refer to this distal region as the UR1 C-terminal segment (UR1cts).

### UR1 is not required for BIK1-induced RBOHD activation in HEK293T cells

UR1 displayed an overall negative regulatory role in Ca²⁺-dependent activation of RBOHD in HEK293T cells. However, RBOHD is also activated by phosphorylation. To determine whether UR1 contributes to phosphorylation-driven activation, or to a combined effect of phosphorylation and Ca²⁺, we co-transfected the 3x-FLAG-RBOHD deletion variants with BIK1, a kinase that phosphorylates RBOHD upon flg22 perception at multiple sites within UR1 and UR2. Expression levels were comparable across constructs (Fig. S5D,E). ROS production was then monitored in HEK293T cells before and after ionomycin treatment.

Before ionomycin treatment, all tested truncation constructs (DΔ60, DΔ69, DΔ83, DΔ100, DΔ121, DΔ134, DΔ155, DΔ170, DΔ181 and DΔ253) displayed similar BIK1-induced activity in basal cytoplasmic Ca²⁺ concentration compared to the full-length RBOHD (Fig. 3E,F). After ionomycin treatment, DΔ60, DΔ69, DΔ83, DΔ100, DΔ121, DΔ134, DΔ155 showed similar activity compared to full length RBOHD, while DΔ181 and DΔ253 showed a drastic reduction. On the other hand, DΔ170 exhibited higher activity after ionomycin treatment, highlighting again the inhibitory function of the segment 155-170 in contrasting the activation caused by Ca²⁺ binding to the EFD (Fig. 3F).

These results suggest that UR1 is not required for BIK1-mediated activation of RBOHD at basal cytoplasmic Ca²⁺ concentrations in HEK293T cells. At elevated cytoplasmic Ca²⁺, however, the 155–170 segment partially opposes RBOHD activation. Given that BIK1-mediated activation is largely maintained in the absence of UR1, and that BIK1 was found inefficient to phosphorylate the C-terminus of RBOHD in HEK293T cells (Köster *et al*, 2025), it is plausible that the activation of RBOHD by BIK1 is mainly driven by phosphorylation of UR2 at three serine residues (S339, S343, and S347), which have previously been identified to be phosphorylated in a BIK1-dependent manner *in vivo* (Li *et al*, 2014; Kadota *et al*, 2014).

### Functional coupling of RBOHH UR1 and the EFD enhances Ca²^⁺^-driven ROS production in HEK293T cells

Previous studies highlighted that RBOHH produces approximately tenfold more ROS than RBOHD in response to Ca²⁺ alone in HEK293T cells (Kaya *et al*, 2019), suggesting that certain features of the RBOHH regulatory architecture are specialized to maximize Ca²⁺-dependent catalytic output compared to RBOHD. Given the contrasting effects of UR1 truncation in RBOHD and RBOHH, particularly near the C-terminal end of UR1, we next asked whether transferring different portions of the RBOHH N-terminal regulatory region could enhance Ca²⁺-dependent activation of RBOHD. For this analysis, we distinguished the main body of UR1 (RBOHD residues 1–161; RBOHH residues 1–100; hereafter UR1-main) from the UR1 C-terminal segment (UR1cts), the short region immediately upstream of the EFD.

We generated chimeric RBOHD variants in which UR1-main alone (χ1), UR1cts together with the EFD (χ2), or the complete UR1 together with the EFD (χ3) were replaced with the corresponding regions from RBOHH (Fig. 3G). All constructs showed similar expression levels in HEK293T cells (Fig. S5F). ROS production was assayed following ionomycin-induced Ca²⁺ influx. Compared with full-length RBOHD, χ1 exhibited a modest increase in activity, χ2 displayed reduced activity, and χ3 showed a strong enhancement of Ca²⁺-induced ROS production (Fig. 3H).

Together, these results suggest that the elevated Ca²⁺-induced activity associated with RBOHH cannot be transferred to RBOHD by UR1-main or by the UR1cts–EFD combination alone. The modest increase observed with χ1 and the reduced activity of χ2 indicate that these partial substitutions are insufficient to confer RBOHH-like activity. In contrast, replacing the complete UR1 with the EFD in χ3 substantially enhanced ROS production, suggesting that UR1 and the EFD function cooperatively in RBOHH to promote strong Ca²⁺-dependent activation in HEK293T cells.

### The RBOHD EFD is required for full BIK1-dependent activation in HEK293T cells

To determine whether introducing regulatory elements from RBOHH into RBOHD also altered phosphorylation-dependent activation via UR2, we co-expressed full-length RBOHD and the RBOHD–RBOHH chimeric constructs with BIK1 in HEK293T cells. Protein levels were similar across all constructs (Fig. S5G). ROS production was measured before and after ionomycin treatment.

Under basal Ca²⁺ conditions, with phosphorylation as the primary driver of activation, wild-type RBOHD and the χ1 chimera (containing the UR1-main region of RBOHH but retaining the RBOHD EFD) exhibited the highest ROS-producing activity (Fig. 3I). By contrast, all constructs containing the EFD from RBOHH (χ2, and χ3) showed markedly reduced ROS production. Following ionomycin treatment, constructs harboring the EFD of RBOHD (WT and χ1) still exhibited higher ROS-producing activity compared to those containing the EFD of RBOHH (Fig. 3I). Together, these results indicate that the RBOHD EFD is required for efficient BIK1-dependent activation in HEK293T cells. The strong reduction in activity observed in both χ2 and χ3, which contain the RBOHH EFD, further suggests that the RBOHD EFD is functionally coupled to phosphorylation-dependent regulation through UR2.

### The RBOHH UR1–EFD module does not fully support flg22-triggered RBOHD activation *in planta*

To determine whether the RBOHH UR1–EFD module could functionally replace the corresponding region of RBOHD during immune signaling, we generated Arabidopsis lines expressing either 3×FLAG–RBOHD or the χ3 chimera under the strong constitutive 35S promoter, with comparable expression levels (Fig. S5H). Leaf discs were treated with flg22, which triggers signaling events including Ca²⁺ influx and phosphorylation of RBOHD by kinases such as BIK1. Although both RBOHD- and χ3-expressing plants produced a characteristic transient ROS burst, χ3 lines exhibited substantially reduced ROS production compared with plants expressing wild-type RBOHD (Fig. 3J). These results suggest that the RBOHH UR1–EFD module cannot fully substitute for the native RBOHD UR1–EFD configuration during flg22-triggered ROS production *in planta*. Notably, the enhanced Ca²⁺-induced activity that the RBOHH UR1–EFD module appears to confer based on the HEK293T cell data was insufficient to compensate for the loss of the native RBOHD module during immune signaling.

### Structural modeling suggests that phosphorylation of S339, S343, and S347 stabilizes the α-helical conformation of RBOHD UR2

Our previous assays indicated that the EFD of RBOHD functionally cooperates with BIK1-dependent phosphorylation of the UR2 region (residues 329–357) to promote enzymatic activation. This is consistent with *in planta* data showing that phosphorylation of UR2 is essential for immune-induced ROS production as the triple non-phosphorylatable variant of RBOHD S339A/S343A/S347A exhibits nearly abolished flg22-triggered ROS (Li *et al*, 2021). However, the molecular mechanism by which phosphorylation of these three serine residues enhances catalytic function has so far not been addressed in detail.

To investigate this, we examined the AlphaFold3-predicted structure of RBOHD and evaluated structural aspects of the UR2 region. Within UR2, AlphaFold3 predicts an α-helix (residues 339 to 349) that contains all three phosphorylatable serines (S339, S343 and S347). However, this α-helix is predicted with a pLDDT score below 50 (Fig. 4A), indicating low confidence. Next, we predicted a structure in which S339, S343, and S347 were phosphorylated. Phosphorylation significantly increased the pLDDT score of the α-helix to the values up to 70 (Fig. 4A).

**Figure 4.**
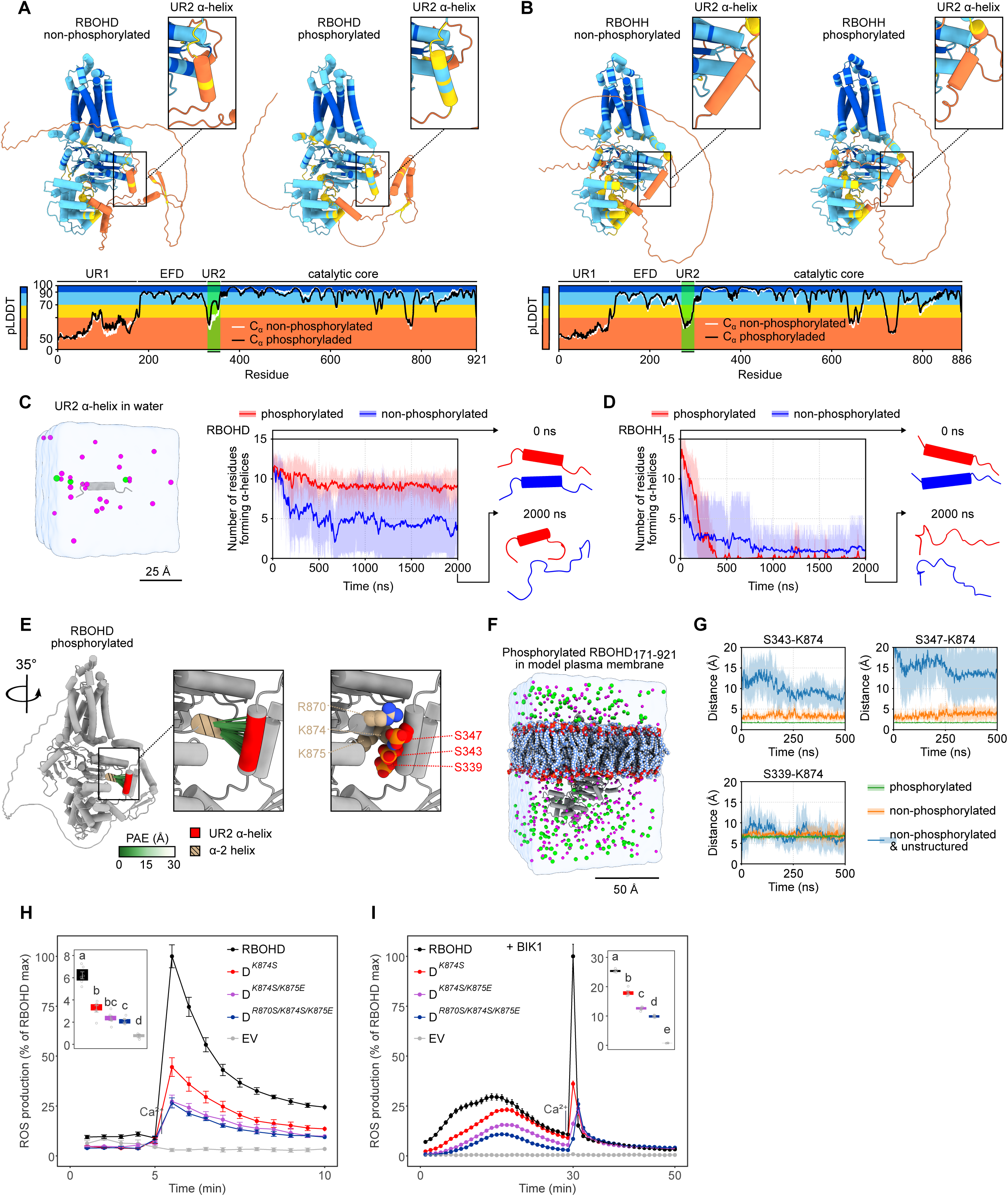
**Structural and functional analysis of the UR2–**α**-2 interaction in RBOHD. (A)** AlphaFold3 predictions of non-phosphorylated RBOHD and RBOHD phosphorylated at S339, S343, and S347. Predicted local distance difference test (pLDDT) scores are shown for each residue, and RBOH domains are annotated above the graph. Blue and light blue indicate high confidence, whereas yellow to red indicates lower confidence. **(B)** AlphaFold3 predictions of non-phosphorylated RBOHH and S283, T285, and T287 phosphorylated RBOHH. **(C)** Molecular dynamics simulations in water of the RBOHD UR2 α-helix containing S339, S343, and S347 in phosphorylated and non-phosphorylated states. Simulations were performed in five independent repeats. Plots show the Gaussian-smoothed average (solid line) with the non-smoothed standard deviation shown transparently. Snapshots of the initial state (0 ns) and one representative final state (2000 ns) are shown for each repeat. **(D)** Molecular dynamics simulations in water of the RBOHH UR2 α-helix containing S283, T285, and T287 in phosphorylated and non-phosphorylated states. **(E)** Position of the phosphorylated UR2 α-helix (residues 339–349) relative to the α-2 helix (869–880) in a AlphaFold3 model of RBOHD. Predicted contacts within 8 Å are colored by Predicted Alignment Error (PAE) (green, high confidence; white, low confidence). Atoms are shown as spheres in the right panel. **(F)** Snapshot from the beginning of the molecular dynamics simulation of RBOHD 171–921 in a model plant plasma membrane**. (G)** Molecular dynamics simulations of RBOHD 171–921 in a model plant plasma membrane. Distances in Å between S339–K874, S343–K874, and S347–K874 over 500 ns. Mean shown as a solid line with standard deviation shown transparently. **(H)** Ca²⁺-induced ROS production in HEK293T cells expressing 3×FLAG–RBOHD or helix α-2 point mutants (K874S, K874S/K875E, R870S/K874S/K875E). ROS measurements were performed as in Fig. 3. **(I)** BIK1-induced and BIK1 + Ca²⁺-induced ROS production in HEK293T cells expressing 3×FLAG–RBOHD or helix α-2 point mutants (K874S, K874S/K875E, R870S/K874S/K875E). ROS measurements were performed as in Fig. 3.

To further test the structural stability of a phosphorylated or unphosphorylated α-helix, we performed all-atom molecular dynamics simulations of the α-helix. The phosphorylated α-helix maintained its α-helical secondary structure, whereas unphosphorylated α-helix gradually transitioned into an unstructured loop (Fig. 4C). These results suggest that phosphorylation at S339, S343, and S347 stabilizes an α-helical conformation within the UR2 region in RBOHD.

In RBOHH, the region homologous to RBOHD UR2 α-helix similarly adopts a α-helical conformation and is predicted with a pLDDT score below 50 in non-phosphorylated state (Fig. 4B). Phosphorylation of RBOHH UR2 has so far not been experimentally observed in RBOHH, however, the region contains three potential phosphorylation sites (S283, T285, and T287). To assess whether phosphorylation could influence the structural behavior of this region in a manner similar to RBOHD, we predicted S283, T285, and T287-phosphorylated RBOHH. In contrast to RBOHD, phosphorylation did not increase pLDDT of the α-helix (Fig. 4B) and during molecular dynamics simulations, both phosphorylated and non-phosphorylated α-helix transitioned into unstructured loops (Fig. 4D). This suggests UR2 in RBOHH does not adopt the α-helical conformation and functions differently than in RBOHD.

### Structural modeling suggests that phosphorylation of RBOHD UR2 strengthens its interaction with helix α-2 in the C-terminus

When evaluating the positioning of UR2 within predicted RBOH structures, we observed that UR2 helix localizes in close proximity to helix α-2 (α-2) in both RBOHD and RBOHH (Fig. 1A). α-2 is located within the catalytic core and is highly consistent in length and relatively conserved in sequence across RBOH subfamilies (Fig. 1G). To further assess the arrangement between UR2 and α-2, we inspected the PAE of the contacts between the UR2 and α-2 helices within 8 Å, a threshold commonly used to define interacting residues. In RBOHD, the positioning of the UR2 helix towards α-2 is predicted with high confidence, as contacts within the 8 Å distance show low PAE values (Fig. S6A). In contrast, the corresponding contacts in RBOHH show higher PAE (Fig. S6B), consistent with the overall high PAE for the position of EFD towards the catalytic core (Fig. S1D).

Given the reliability of the UR2 helix positioning toward α-2, we further examined the interaction interface in RBOHD. Three positively charged residues (R870, K874 and K875) are present in α-2 in RBOHD. These basic residues in RBOHD likely promote electrostatic interactions with phosphorylatable serines within the UR2 helix. In RBOHH, the absence of corresponding positive charges in α-2 results in less stable contacts, reflected in the lower prediction confidence. Consistent with this model, introducing the phosphate groups to S339, S343, and S347 in UR2 of RBOHD further increases the prediction confidence of UR2–α-2 contacts (Fig. 4E).

To get more insight into the impact of S339, S343 and S347-phosphorylation on the interaction between RBOHD UR2 helix and α-2, we performed all-atomistic molecular dynamics simulations of a RBOHD171-921 embedded into a model plant plasma membrane (Fig. 4F). A truncated version of the protein was employed because AlphaFold does not reliably predict the positioning of UR1 relative to the rest of the protein. Additionally, an extended UR1 would hinder simulation convergence. Over the time course of the simulation, we tracked the distances between each positively charged residue of α-2 (R870, K874, K875) and the phosphorylated or non-phosphorylated serines of UR2 (S339, S343, S347). For the non-phosphorylated RBOHD171-921 two versions of UR2 were tested: one containing a helical UR2 region and another in which the UR2 (residues 335 to 354) was modeled as an unstructured loop, as we showed the unphosphorylated UR2 helix become unstructured over time. We observed that phosphorylation keeps UR2 in close contact with α-2. The interaction is primarily mediated by S343 and S347 interacting with K874 (Fig. 4G), supported by the interaction of S347 with R870 and S339 with K875 (Fig. S6C,D).

Collectively, these analyses support a model in which phosphorylation of S339, S343, and S347 stabilizes a helical conformation within UR2 and, by introducing the negative charge, strengthens the interaction of UR2 with α-2 located within the RBOHD catalytic core.

### Positively charged residues in α-2 contribute to Ca²^⁺^- and BIK1-induced RBOHD activation in HEK293T cells

Because the proposed interface between UR2 and α-2 in RBOHD depends on electrostatic attraction between negatively charged phosphoserines in UR2 and positively charged amino acids in α-2, we next examined whether disrupting the positive charges would compromise RBOHD function. Thus, we generated several RBOHD point mutants: K874S, K874S/K875E, and R870S/K874S/K875E. Residues K874 and K875 were substituted with serine (S) and glutamate (E), respectively, to match homologous residues in RBOHH (Fig. S6E). Additionally, residue R870, which lacks a corresponding negatively charged residue in RBOHH, was replaced with serine (S) to completely remove positive charges from α-2 in RBOHD.

RBOHD K874S, K874S/K875E, and R870S/K874S/K875E variants were expressed in HEK cells with or without BIK1. Protein levels were similar across all variants (Fig. S5I,J). ROS-producing activity was measured before and after ionomycin treatment. Following ionomycin treatment, the RBOHD K874S mutant exhibited reduced activity compared to wild type RBOHD (Fig. 4H). Additional substitution in K874S/K875E led to a further reduction in activity, while no additional loss was observed upon further mutation R870S in the R870S/K874S/K875E variant (Fig. 4H). When co-expressed with BIK1, all RBOHD variants (K874S, K874S/K875E, and R870S/K874S/K875E) showed a progressive reduction in activity before ionomycin treatment, with the triple point mutant displaying the lowest activity (Fig. 4I). After ionomycin-induced calcium elevation, the mutants exhibited activity profiles similar to those observed without BIK1 co-expression (Fig. 4I).

Together, these results indicate that positively charged residues in α-2 are required for full activation of RBOHD under both Ca²⁺-induced and BIK1-induced conditions. The unexpected effect of the α-2 mutations on Ca²⁺-dependent activation suggests that Ca²⁺ binding and BIK1-dependent phosphorylation may converge on a shared mechanism involving stabilization of the UR2–α-2 interface to promote catalysis. Notably, α-2 in RBOHH lacks the basic residues present in RBOHD: the positions corresponding to K874 and K875 in RBOHD are occupied by serine and glutamate in RBOHH, creating a neutral or negatively charged surface. This configuration is incompatible with the electrostatic interactions proposed to stabilize the UR2–α-2 interface in RBOHD, further suggesting that the unstructured regulatory regions of RBOHD and RBOHH have undergone functional diversification.

## Discussion

ROS generated by RBOHs are central to plant physiology, mediating immune defense, abiotic stress responses, and polarized cell growth. Ca²⁺-binding and phosphorylation are well-established activating inputs for RBOHs. However, how the unstructured regions contribute to Ca²⁺- and phosphorylation-dependent RBOH activation remains largely unexplored. Here, we investigated two major unstructured regulatory regions, UR1 and UR2, in Arabidopsis RBOHD and RBOHH, two evolutionarily distant RBOHs with distinct physiological functions. RBOHD is broadly expressed and participates in multiple stress-related and developmental processes, whereas RBOHH expression is largely restricted to pollen tubes, where oscillatory Ca²⁺ signals accompany polarized growth. Our results support distinct regulatory roles for the unstructured regions of RBOHD and RBOHH and suggest that their divergence contributes to differences in Ca²⁺- and phosphorylation-dependent activation.

UR1 is located at the very N-terminus of RBOHs and directly connects to the EFD. This region varies in both length and amino acid composition across RBOH paralogs throughout the plant lineage. Our truncation analyses revealed contrasting effects of UR1 removal in RBOHD and RBOHH in HEK293T cells. In RBOHD, complete UR1 removal increased ROS production under elevated Ca²⁺ conditions, suggesting an overall inhibitory role for this region. The strongest inhibitory effect was associated with the C-terminal portion of UR1 (residues 155–170). The truncation series also suggested additional regulatory elements within UR1, including residues 121–134, whose removal restored activity reduced by preceding truncations. By contrast, progressive truncation of RBOHH UR1 resulted in a substantial loss of Ca²⁺-induced ROS production. Together, these observations suggest that UR1 contributes differently to Ca²⁺-dependent regulation of RBOHD and RBOHH: UR1 appears to restrain activation in RBOHD, whereas it positively contributes to activation in RBOHH. The molecular basis underlying these contrasting effects remains to be determined.

RBOHD and RBOHH also differ markedly in their intrinsic Ca²⁺ responsiveness, with RBOHH previously reported to produce approximately tenfold more ROS than RBOHD following Ca²⁺ elevation in HEK293T cells. Our domain-swap experiments suggest that the enhanced Ca²⁺ responsiveness associated with RBOHH depends on coordinated features of its UR1–EFD module. Replacement of the complete UR1 together with the EFD from RBOHH in RBOHD substantially enhanced Ca²⁺-induced ROS production, whereas substitution of only portions of this module resulted in relatively minor changes in activity. However, in HEK293T cells, RBOHD chimeric variants containing the EFD of RBOHH displayed impaired activation by BIK1, which phosphorylates S339, S343, and S347 in UR2 of RBOHD (Li *et al*, 2014; Kadota *et al*, 2014). The chimeric variant containing both UR1 and the EFD from RBOHH also exhibited reduced ROS production *in planta* following flg22 perception compared with wild-type RBOHD. Together, these results suggest that the EFD of RBOHD may have evolved to operate in concert with UR2 to support phosphorylation-dependent activation, whereas in RBOHH the EFD appears to be functionally coupled with UR1 to promote stronger Ca²⁺-dependent activation.

Besides UR1 and the EFD, our analyses suggest that functional specialization may also extend to UR2. Phosphorylation of S339, S343, and S347 within RBOHD UR2 has been repeatedly shown to be crucial for immune-induced activation (Kadota *et al*., 2015; Li *et al*., 2021). However, how phosphorylation of these residues is translated into increased enzymatic activity remains unclear, and whether the corresponding UR2 region of RBOHH could support a similar regulatory mechanism is unknown.

Our structural analyses suggest a possible mechanism. Molecular dynamics simulations indicated that phosphorylation of S339, S343, and S347 favors stabilization of an α-helical conformation within RBOHD UR2, positioning the three phosphoserines in close proximity to the positively charged residues K874 and K875 in α-2 of the C-terminus. This arrangement could favor electrostatic interactions between UR2 and α-2. Consistent with this model, disrupting the positive charge at this predicted interface by substituting K874 and K875 with the corresponding residues from RBOHH, serine and glutamate (K874S/K875E), resulted in a strong reduction in BIK1-induced ROS production in HEK293T cells. Interestingly, disruption of the same predicted interface also reduced Ca²⁺-induced activity, suggesting that the UR2–α-2 interaction could contribute to both phosphorylation- and Ca²⁺-dependent activation.

Although we did not experimentally investigate the structural basis of this effect, one possibility is that stabilization of the UR2–α-2 interface reduces the mobility of the catalytic region, of which α-2 is part, thereby favoring electron transfer. A conceptually similar mechanism has been proposed for mammalian DUOX1, where Ca²⁺-dependent interactions between the EF-hand module and the dehydrogenase domain reduce its mobility and promote catalytic activity (Wu *et al*, 2021). Our analyses suggest that the same mechanism is less likely to operate in RBOHH. *In silico* phosphorylation of the corresponding potential phosphorylation sites in RBOHH UR2 (S283, T285, and T287) did not promote stabilization of the predicted α-helix. In addition, α-2 in RBOHH lacks the positive charges corresponding to K874 and K875 in RBOHD, making an equivalent electrostatic interaction less favorable. Together, these differences suggest distinct regulatory roles for UR2 in RBOHD and RBOHH.

The contrasting regulatory architectures identified in this study may relate to the distinct physiological environments in which RBOHD and RBOHH operate. Pollen tubes, where RBOHH is specifically expressed, feature dynamic oscillations in cytosolic Ca²⁺ at the growing tip (Kaya *et al*, 2014). The proposed functional coupling between UR1 and the EFD in RBOHH may enhance Ca²⁺ responsiveness and could therefore be well suited to the rapid Ca²⁺ dynamics associated with pollen tube growth. In contrast, the broadly expressed RBOHD is central to responses to biotic and abiotic stimuli, plant development, and systemic ROS signaling. The inhibitory role of UR1 in RBOHD may help limit activation by physiological background Ca²⁺ fluctuations, thereby restricting unnecessary ROS production. At the same time, the apparent functional coupling between the EFD and UR2 could favor stronger phosphorylation-dependent control and may contribute to sustaining RBOHD activity over longer timescales during immune signaling. Taken together, our results suggest that evolutionary diversification of regulatory regions has contributed to tuning RBOH activation to distinct physiological contexts.

## Author Contributions

M.C., R.P., and M.W. designed research; M.C. performed HEK293T cells experiments; M.N. performed structural modeling and molecular dynamics simulations; A.P. generated transgenic plants and helped establish the HEK293T assay; A.Z. performed ROS measurements in plants; P.P. and T.K.P. contributed to experimental design and interpretation of preliminary experiments that informed subsequent experimental directions; M.P. performed phylogenetic analyses; S.K., R.P., and M.W. supervised research; and M.C., P.P., R.P., M.P., and M.W. secured funding.

## Acknowledgments

We dedicate this article to the memory of Jaakko Kangasjärvi - advisor, mentor and friend. We thank Mr. Jan Kadlec (Growth Facility, Biology Centre CAS Core Facilities) for technical assistance, Dr. Cezary Waszczak (University of Helsinki) for providing molecular biology materials, and Dr. Yaseen Mottiar (University of Ottawa) for helping secure part of the funding for this research. Computational resources were provided by the e-INFRA CZ project (ID: 90254), supported by the Ministry of Education, Youth and Sports of the Czech Republic.

## Data Availability Statement

Sequence datasets and alignments used for the phylogenetic analyses are provided as Supplementary Datasets S1–S5. Other data supporting the findings of this study are contained within the article and its Supporting Information or are available from the corresponding authors upon reasonable request.

## AI Generative Statement

Generative AI (ChatGPT, OpenAI; GPT-5.6 Sol) was used to improve sentence structure, clarity, and flow. All AI-assisted content was reviewed and edited by the authors.

## Supporting Information

Additional Supporting Information is available with this article.

**Supplementary Methods** Extended methods for phylogenetic analyses and molecular dynamics simulations.

**Figures S1–S6.** Supplementary structural, sequence, phylogenetic, protein-expression, and molecular dynamics analyses.

**Datasets S1–S5.** Sequence datasets and alignments used for phylogenetic analyses.

