## Supplementary Information for "Unstructured regions differentially modulate the activation of RBOHD and RBOHH"

**This PDF includes**

Supplementary Methods

Figures S1–S6

Supplementary Datasets S1–S5 are provided separately.

### Supplementary Methods

#### Phylogenetic Analysis

Homologous sequences were identified using BLASTP and TBLASTN searches against the NCBI, Phytozome, and Phycocosm databases using Arabidopsis RBOHD, RBOHH, and FRO1 (AT1G01590) as queries. Human NOX2, NOX5, DUOX1, and yeast FRE1 sequences were also used to aid identification of related NADPH oxidase family members. Protein sequences were aligned using MAFFT (E-INS-I algorithm) implemented in Jalview, and gap-rich and poorly conserved regions were manually trimmed.

For the global phylogenetic analysis, a dataset of 304 sequences and 414 aligned positions was analyzed using IQ-TREE with the Q.pfam+R10 substitution model and ten free-rate categories. For the embryophyte RBOH analysis, a dataset of 479 sequences and 556 aligned positions was analyzed using IQ-TREE with the Q.PLANT+R9 model and nine free-rate categories. Ultrafast bootstrap analysis (10,000 replicates) was performed to assess branch support.

Bayesian phylogenetic inference of the embryophyte RBOH dataset was performed using MrBayes with a mixed amino acid substitution model. Two independent runs with four chains were performed for 2,000,000 generations, sampling trees every 100 generations. Convergence was reached after approximately 500,000 generations, which were discarded as burn-in. Trees were visualized and analyzed using TreeViewer. The complete sequence list and alignments are provided in Supplementary Datasets S1–S5.

#### Structural Modelling and Molecular Dynamics Simulations

For simulations of the UR2 helix in water, the UR2 helix, including several adjacent residues (RBOHD 335–354 and RBOHH 275–294), was placed in a cubic simulation box (6.5 × 6.5 × 6.5 $nm^3$). For simulations of the phosphorylated UR2 helix, phosphorylation was introduced at S339, S343, and S347 of RBOHD or S283, T285, and T287 of RBOHH. Systems were solvated with water containing 0.15 M KCl and neutralized.

Energy minimization was carried out using the steepest-descent algorithm for up to 5000 steps, followed by a 125 ps equilibration. During both minimization and equilibration, position restraints were applied to protein backbone and side-chain atoms. A 2000 ns production run was subsequently performed in the NPT ensemble with a 2 fs time step. Pressure was maintained at 1 bar using the Parrinello–Rahman barostat (coupling constant 5.0 ps; compressibility $4.5 \times 10^{-4}$). Temperature was maintained at 303.15 K using the v-rescale thermostat (coupling constant 1.0 ps). Secondary structure was subsequently analyzed using the GROMACS built-in tool gmx dssp, and the number of amino acid residues adopting an α-helical conformation was plotted over time.

For simulations of phosphorylated and non-phosphorylated RBOHD171–921 in a model plant plasma membrane, the protein was placed in a 12 × 12 × 16.5 $nm^3$ simulation box and embedded in a lipid bilayer with the following composition: outer leaflet POPC:POPE (50:50), inner leaflet POPC:POPE:POPS:POPA:POPI4P:POPI(4,5)P2 (37:37:10:10:5:1). The charge of the POPA headgroup was modified to −2. Phosphorylation of RBOHD171–921 was introduced at residues S339, S343, and S347.

Two variants of non-phosphorylated RBOHD171–921 were simulated: one containing a helical UR2 region and another in which UR2 (residues 335–354) was replaced by a flexible loop generated using MODELLER. Systems were solvated with water containing 0.15 M KCl and neutralized. Energy minimization was carried out using the steepest-descent algorithm for up to 5000 steps, followed by a 1875 ps equilibration. During minimization and equilibration, position restraints were applied to protein backbone and side-chain atoms, lipid phosphate atoms, and dihedral angles. A 500 ns production run was subsequently performed under the same conditions used for the UR2 helix simulations. Minimum distances between S339, S343, and S347 and R870, K874, and K875 were calculated using the GROMACS built-in tool gmx mindist.

### Figures

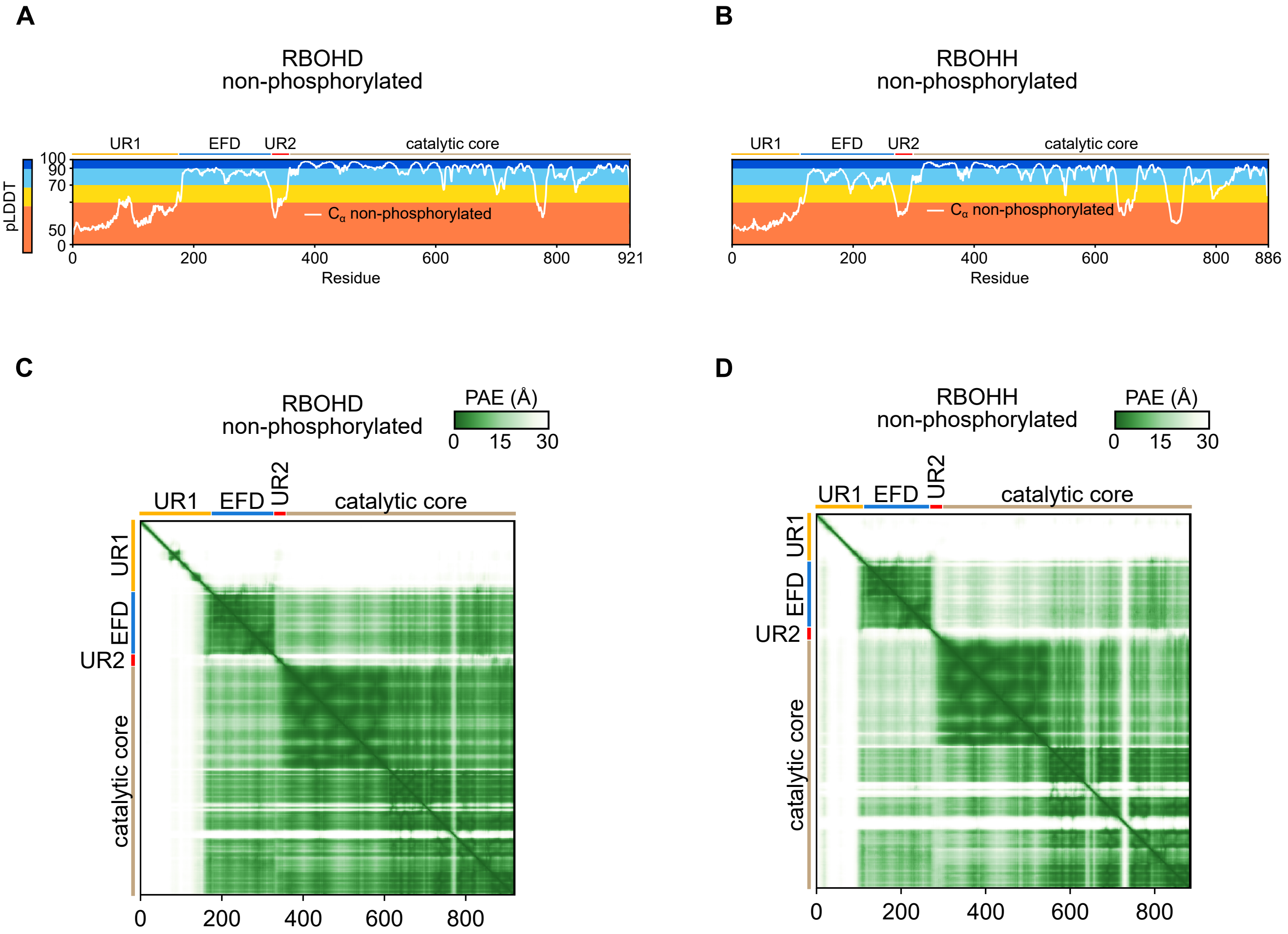


**Fig. S1 (A)** AlphaFold3 predictions of non-phosphorylated RBOHD. Predicted local distance difference test (pLDDT) scores are shown for each residue, with RBOH domains annotated above the graph. Blue and light blue indicate high confidence, whereas yellow to red indicate lower confidence. **(B)** AlphaFold3 predictions of non-phosphorylated RBOHH. pLDDT scores are shown as in panel A. **(C)** Predicted Alignment Error (PAE) of AlphaFold3 predictions of non-phosphorylated RBOHD. Green indicates high confidence, whereas white indicates low confidence. **(D)** Predicted Alignment Error (PAE) of AlphaFold3 predictions of non-phosphorylated RBOHH.

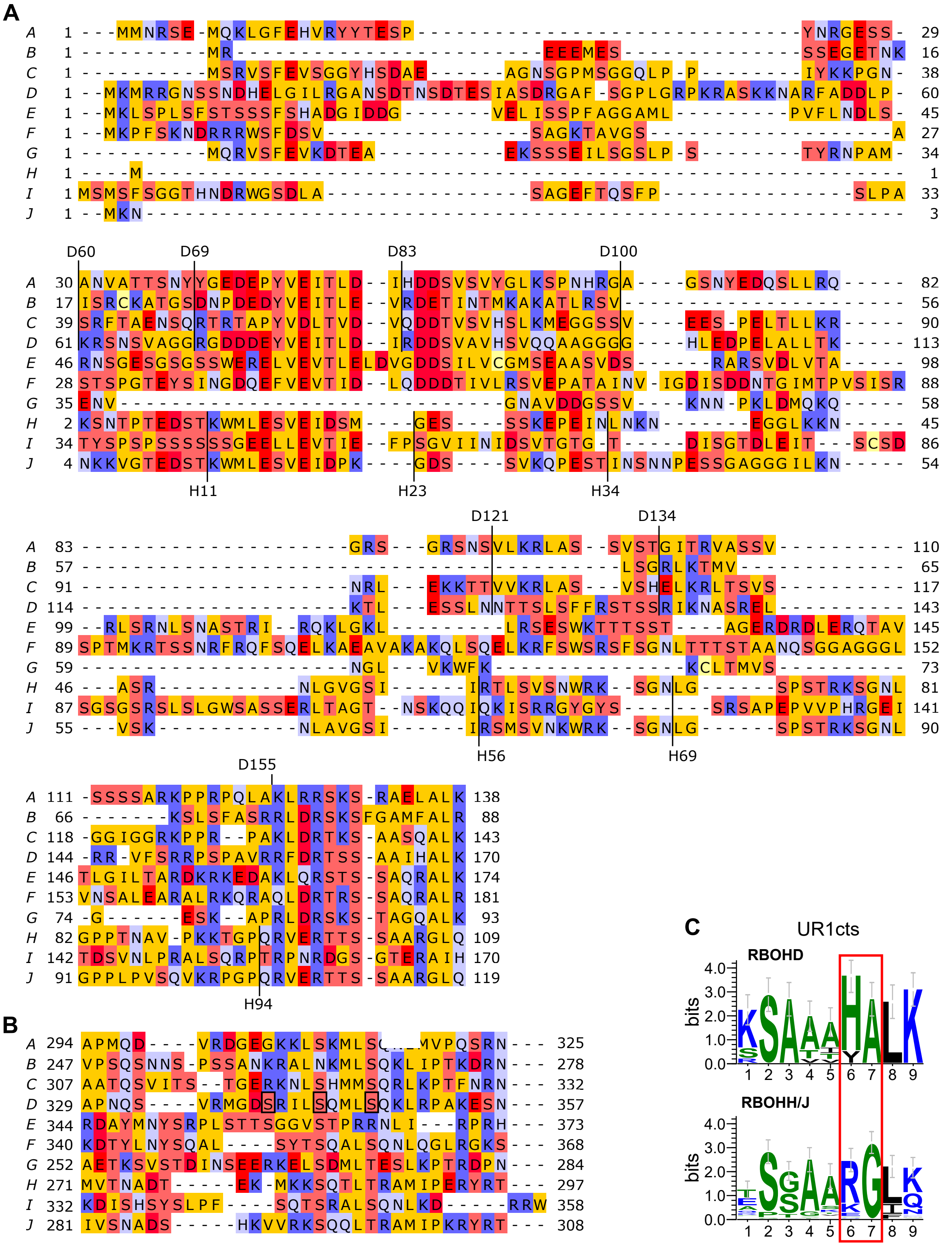


**Fig. S2 (A)** Alignment of the unstructured region 1 (UR1) of the ten Arabidopsis RBOH proteins. Residues used as starting points for the deletion constructs of RBOHD are indicated by a black line and annotated above the alignment; the corresponding residues in RBOHH are annotated below. Alignments were visualized and colored in Jalview according to residue chemical properties: hydrophobic (yellow), acidic (red), basic (blue), and polar uncharged (light red or light blue). **(B)** Alignment of the unstructured region 2 (UR2). Residues S339, S343, and S347 of RBOHD are outlined with black boxes. **(C)** Sequence logo of the UR1 C-terminal segment (UR1cts) region from embryophyte RBOHD and RBOHH/J subfamily proteins. Amino acid conservation is shown as relative letter height at each position.

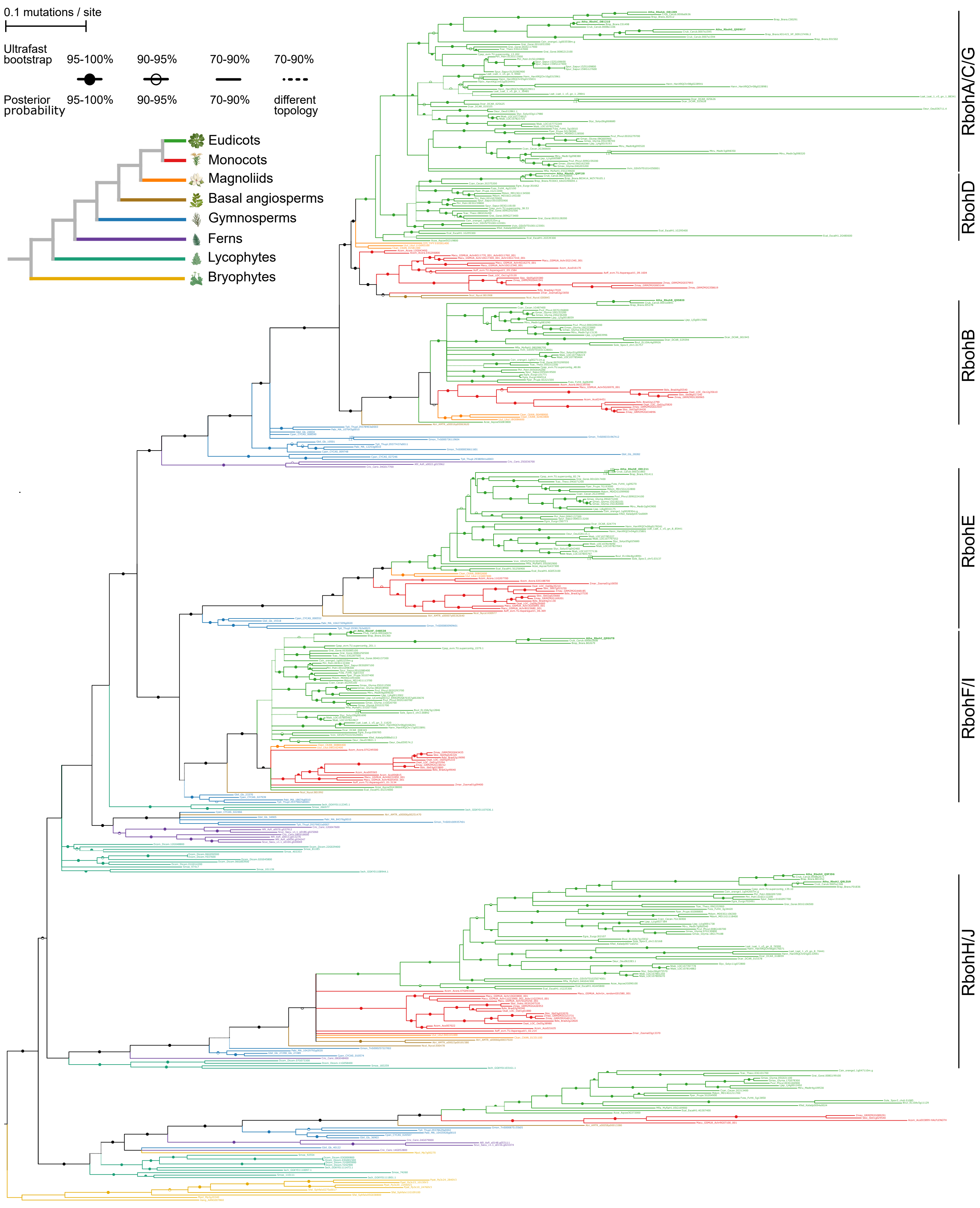
0.1 mutations / site
Ultrafast bootstrap
95-100%
90-95%
70-90%
70-90%
Posterior probability
95-100%
90-95%
70-90%
different topology
Eudicots
Monocots
Magnoliids
Basal angiosperms
Gymnosperms
Ferns
Lycophytes
Bryophytes
RbohA/C/G
RbohD
RbohB
RbohE
RbohF/I
RbohH/J

**Fig. S3.** Maximum-likelihood phylogenetic tree of RBOH protein sequences from representative land-plant species. A largely consistent Bayesian phylogeny was also obtained; clades with differing Bayesian topology are shown with dashed lines. Support for the maximum-likelihood tree was assessed using ultrafast bootstrapping, and posterior probabilities indicate full Bayesian support. Arabidopsis RBOH isoforms are shown in bold. Six robustly supported subfamilies (RbohA/C/G, B, D, E, F/I, and H/J) are indicated. The color code denotes major embryophyte groups, and their evolutionary relationships are shown in the accompanying schematic cladogram. Species abbreviations: Atri – Amborella trichopoda, Acom – Ananas comosus, Aang – Anthoceros angustus, Atha – Arabidopsis thaliana, Aoff – Asparagus officinalis, Afil – Azolla filiculoides, Bvul – Beta vulgaris, Bdis – Brachypodium distachyon, Brap – Brassica rapa, Crub – Capsella rubella, Cpap – Carica papaya, Crich – Ceratopteris richardii, Ccan – Cercis canadensis, Ckan – Cinnamomum kanehirae, Csin – Citrus sinensis, Cpan – Cycas panzhihuaensis, Dcar – Daucus carota, Dcom – Diphasiastrum complanatum, Ecal – Eschscholzia californica, Egra – Eucalyptus grandis, Fves – Fragaria vesca, Gbil – Ginkgo biloba, Gmax – Glycine max, Gmon – Gnetum montanum, Grai – Gossypium raimondii, Hann – Helianthus annuus, iech – Isoetes echinospora, Kfed – Kalanchoe fedtschenkoi, Ls at – Lactuca sativa, Ltul – Liriodendron tulipifera, Ljap – Lotus japonicus, Mdom – Malus domestica, Mpol – Marchantia polymorpha, Mtru – Medicago truncatula, Macu – Musa acuminata, Mfla – Myrothamnus flabellifolia, Ntab – Nicotiana tabacum, Ncol – Nymphaea colorata, Oeur – Olea europaea, Osat – Oryza sativa, Pvul – Phaseolus vulgaris, Pabi – Picea abies, Ppat – Physcomitrella patens, Ptri – Populus trichocarpa, Pper – Prunus persica, Spur – Salix purpurea, Scuc – Salvinia cucullata, Sm oe – Selaginella moellendorffii, Slyc – Solanum lycopersicum, Sbic – Sorghum bicolor, Sfal – Sphagnum fallax, Sole – Spinacia oleracea, Tcac – Theobroma cacao, Tpli – Thuja plicata, Vvin – Vitis vinifera, Zmay – Zea mays, Zmar – Zostera marina.

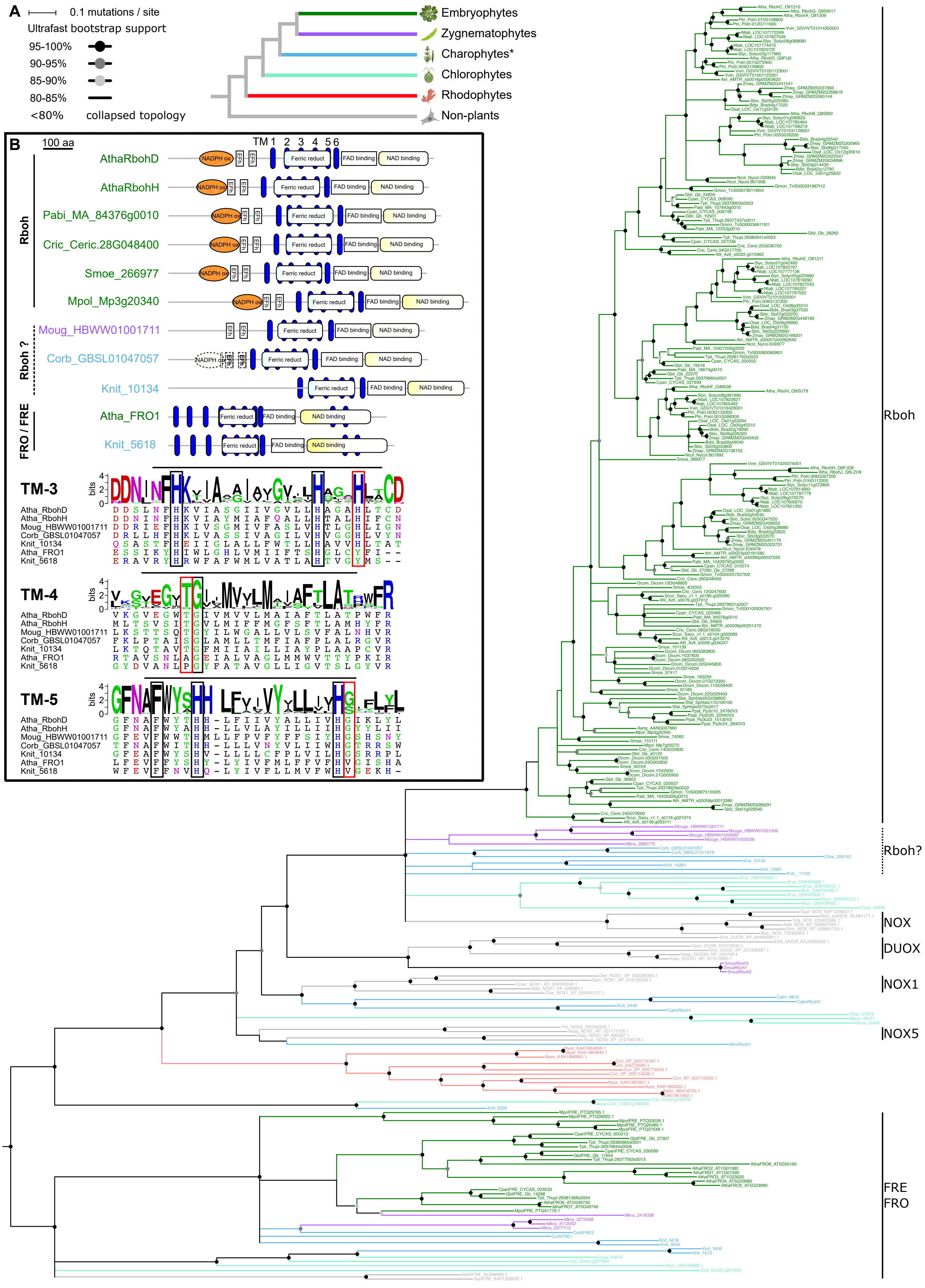
A
0.1 mutations / site
Ultrafast bootstrap support
95-100%
90-95%
85-90%
80-85%
<80%
collapsed topology
Embryophytes
Zygnematophytes
Charophytes*
Chlorophytes
Rhodophytes
Non-plants
B
100 aa
TM 1 2 3 4 5 6
Rboh
AthaRbohD
AthaRbohH
Pabi_MA_84376g0010
Cric_Ceric.28G048400
Smoe_266977
Mpol_Mp3g20340
Rboh ?
Moug_HBWW01001711
Corb_GBSL01047057
Knit_10134
FRO / FRE
Atha_FRO1
Knit_5618
NADPH ox
EFh
Ferric reduct
FAD binding
NAD binding
TM-3
TM-4
TM-5
bits
Atha_RbohD
Atha_RbohH
Moug_HBWW01001711
Corb_GBSL01047057
Knit_10134
Atha_FRO1
Knit_5618
Rboh
Rboh?
NOX
DUOX
NOX1
NOX5
FRE
FRO

**Fig. S4.** RBOH origin within the NADPH oxidase/FRE superfamily. **(A)** Detailed phylogeny of NADPH oxidase– and ferric reductase–like proteins corresponding to Fig. 2B, showing a single, well-supported RBOH clade restricted to streptophyte algae and land plants. Animal NOX/DUOX and plant FRE/FRO families form separate groups, while the evolutionary placement of chlorophyte and sporophyte RBOH-like proteins is scattered. The maximum-likelihood tree is shown with ultrafast bootstrap support indicated by internode circles in different shades of gray. Nodes with support below 80% were collapsed. The color code denotes major Archaeplastida lineages (note that "Charophytes*" represents a polyphyletic grouping of several streptophyte algal clades). Non-plant clades are shown in gray. **(B)** Upper panel: Domain comparisons of representative RBOH and FRO/FRE proteins from embryophytes, zygnematophytes, and streptophyte algae. Canonical RBOHs contain paired N-terminal EF-hand motifs, which are absent from FRO/FRE proteins, whereas both groups retain the ferric-reductase–like transmembrane domain and FAD/NAD(P)H-binding subdomains. Lower panel: Sequence logos for transmembrane segments TM3–TM5, highlighting conserved heme-ligating residues (black boxes) and positions that distinguish NADPH oxidases from ferric reductases (red boxes).

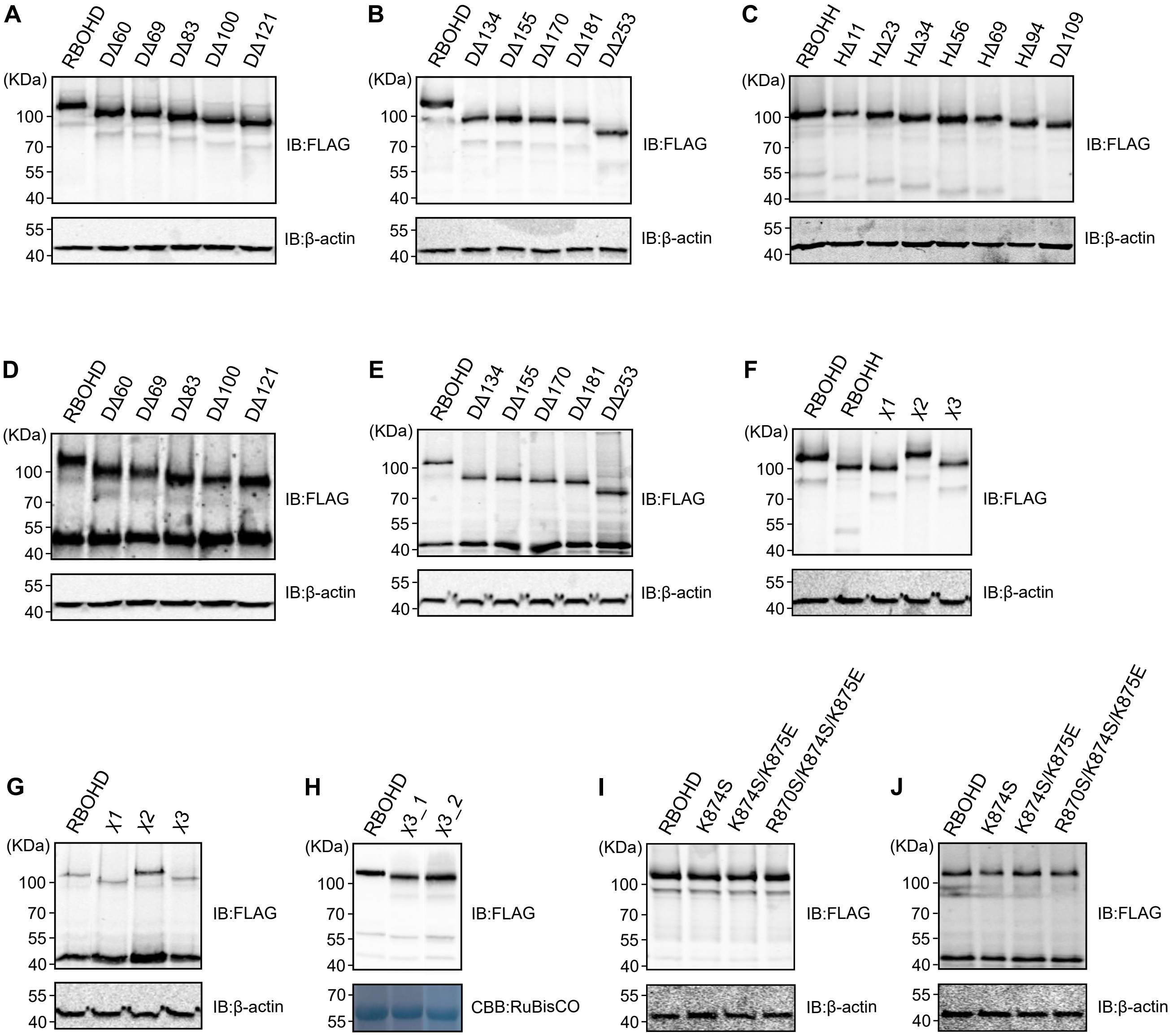


**Fig S5.** Immunoblot analyses. All RBOHD- and RBOHH-based constructs and BIK1 contain an N-terminal 3×FLAG tag. Approximate molecular weights of the expressed proteins are as follows: RBOHD: 106.8 kDa; DΔ1–60: 100.3 kDa; DΔ1–69: 99.6 kDa; DΔ1–83: 97.9 kDa; DΔ1–100: 96.3 kDa; DΔ1–121: 94.0 kDa; DΔ1–134: 92.6 kDa; DΔ1–155: 90.1 kDa; DΔ1–170: 88.4 kDa; DΔ1–181: 87.3 kDa; DΔ1–253: 79.0 kDa; RBOHH: 103.5 kDa; HΔ1–11: 102.4 kDa; HΔ1–23: 101.0 kDa; HΔ1–34: 99.8 kDa; HΔ1–56: 97.6 kDa; HΔ1–69: 96.1 kDa; HΔ1–94: 93.6 kDa; HΔ1–109: 92.0 kDa; χ1: 99.9 kDa; χ2: 107.7 kDa; χ3: 100.9 kDa; K874S: 106.7 kDa; K874S/K875E: 106.7 kDa; R870S/K874S/K875E: 106.7 kDa; BIK1: 46.8 kDa.
β-actin (42 kDa) and RuBisCO subunit (55 kDa) were used as loading controls.

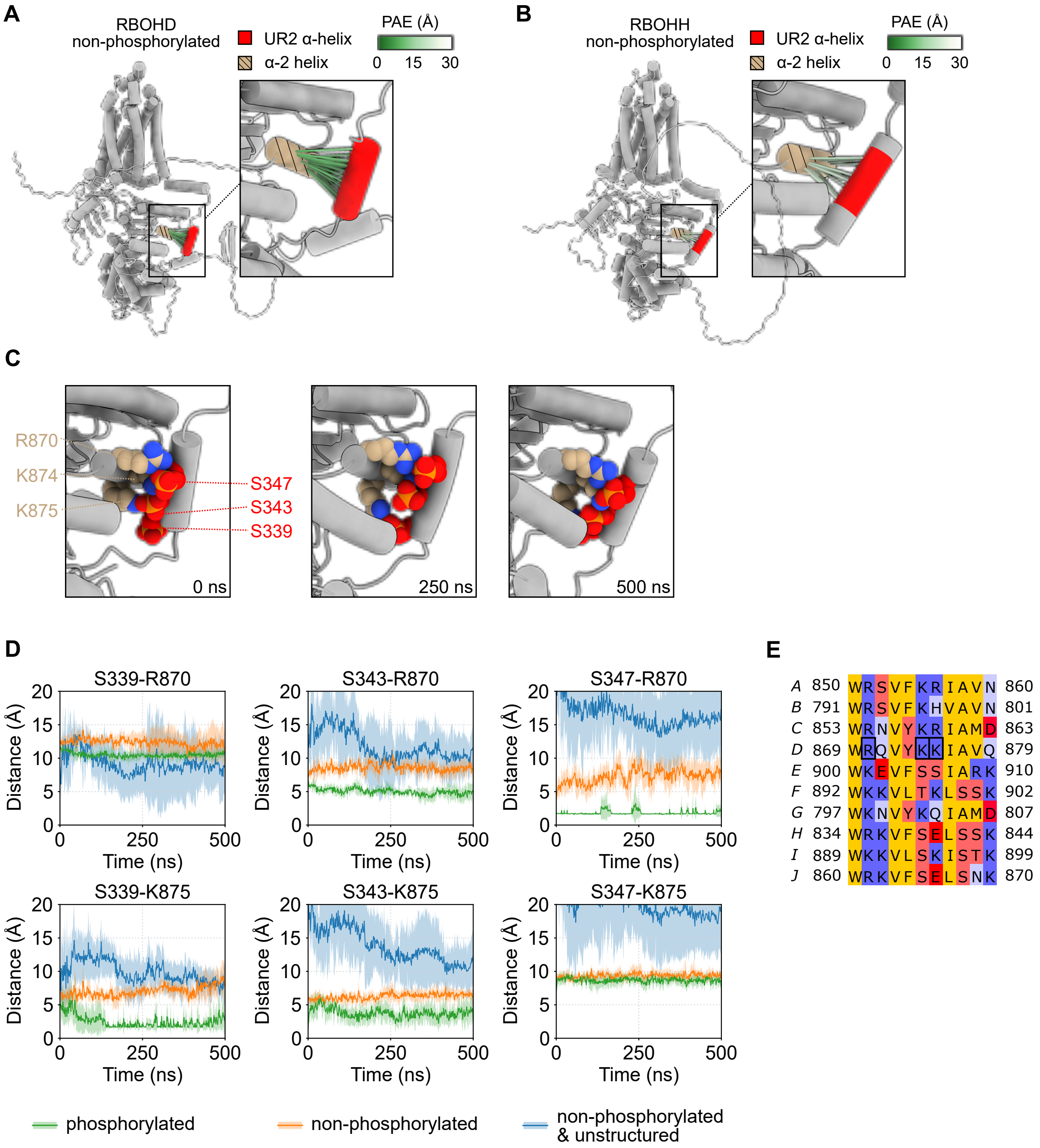


**Fig. S6. (A)** Position of the non-phosphorylated UR2 α-helix relative to the α-2 helix in a AlphaFold3 model of RBOHD. Predicted contacts within 8 Å are colored by Predicted Alignment Error (PAE) (green, high confidence; white, low confidence). **(B)** Position of the non-phosphorylated UR2 α-helix relative to the α-2 helix in a AlphaFold3 model of RBOHH. Predicted contacts within 8 Å are colored by Predicted Alignment Error (PAE) (green, high confidence; white, low confidence). **(C)** Snapshots from molecular dynamics simulations of RBOHD 171–921 in a model plant plasma membrane showing interaction of S339, S343, and S347 of the UR2 α-helix and the positively charged residues R870, K874, and K875 of the α-2 helix 2 at 0 ns, 250 ns and 500 ns. **(D)** Minimum distances for serine–arginine/lysine pairs not shown in Fig. 4G. S347 remains in close proximity to R870 and S339 to K875 over the 500 ns time. The plot displays average (solid line) with a standard deviation shown transparently. **(E)** Alignment of helix α-2 from the catalytic core. Residues R870, K874, and K875 of RBOHD are outlined with black boxes.
